# *Scutellaria baicalensis* extract and baicalein inhibit replication of SARS-CoV-2 and its 3C-like protease *in vitro*

**DOI:** 10.1101/2020.04.10.035824

**Authors:** Hongbo Liu, Fei Ye, Qi Sun, Hao Liang, Chunmei Li, Roujian Lu, Baoying Huang, Wenjie Tan, Luhua Lai

## Abstract

COVID-19 has become a global pandemic that threatens millions of people worldwide. There is an urgent call for developing effective drugs against the virus (SARS-CoV-2) causing this disease. The main protease of SARS-CoV-2, 3C-like protease (3CL^pro^), is highly conserved across coronaviruses and is essential for the maturation process of viral polyprotein. Scutellariae radix (Huangqin in Chinese), the root of *Scutellaria baicalensis* has been widely used in traditional Chinese medicine to treat viral infection related symptoms. The extracts of *S. baicalensis* have exhibited broad spectrum antiviral activities. We studied the anti-SARS-CoV-2 activity of *S. baicalensis* and its ingredient compounds. We found that the ethanol extract of *S. baicalensis* inhibits SARS-CoV-2 3CL^pro^ activity *in vitro* and the replication of SARS-CoV-2 in Vero cells with an EC_50_ of 0.74 μg/ml. Among the major components of *S. baicalensis*, baicalein strongly inhibits SARS-CoV-2 3CL^pro^ activity with an IC_50_ of 0.39 μM. We further identified four baicalein analogue compounds from other herbs that inhibit SARS-CoV-2 3CL^pro^ activity at microM concentration. Our study demonstrates that the extract of *S. baicalensis* has effective anti-SARS-CoV-2 activity and baicalein and analogue compounds are strong SARS-CoV-2 3CL^pro^ inhibitors.

## Introduction

Coronaviruses (CoVs) are single stranded positive-sense RNA viruses that cause severe infections in respiratory, hepatic and various organs in humans and many other animals[1, 2]. Within the 20 years of the 21st century, there are already three outbreaks of CoV-causing global epidemics, including SARS, MERS, and COVID-19. The newly emerged CoV infectious disease (COVID-19) already caused more than 1.5 million confirmed infections and 89 thousands deaths worldwide up to April 9, 2020 (https://www.who.int/emergencies/diseases/novel-coronavirus-2019/situation-reports). There is an urgent call for drug and vaccine research and development against COVID-19.

COVID-19 was confirmed to be caused by a new coronavirus (SARS-CoV-2), whose genome was sequenced in early January 2020[3, 4]. The genomic sequence of SARS-CoV-2 is highly similar to that of SARS-CoV with 79.6% sequence identity [5] and remain stable up to now[6]. However, the sequence identities vary significantly for different viral proteins[7]. For instance, the spike proteins (S-protein) in CoVs are diverse in sequences and even in the host receptors that bind due to the rapid mutations and recombination[8]. Although it has been confirmed that both SARS-CoV and SARS-CoV-2 use ACE2 as receptor and occupy the same binding site, their binding affinities to ACE2 vary due to subtle interface sequence variations[9]. On the contrary, the 3C-like proteases (3CL^pro^) in CoVs are highly conserved. The 3CL^pro^ in SARS-CoV and SARS-CoV-2 share a sequence identity of 96.1 %, making it an ideal target for broad spectrum anti-CoV therapy.

Although many inhibitors have been reported for SARS-CoV and MERS-CoV 3CL^pro^[10–13], unfortunately none of them has entered into clinical trial. Inspired by the previous studies, several covalent inhibitors were experimentally identified to inhibit the 3CL^pro^ activity and viral replication of SARS-CoV-2, and some of the complex crystal structures were solved[14, 15]. In addition, a number of clinically used HIV and HCV protease inhibitors have been proposed as possible cure for COVID-19 [16] and some of them are now processed to clinically trials[17]. Several computational studies proposed potential SARS-CoV-2 3CL^pro^ inhibitors by virtual screening against the crystal or modeled three-dimensional structure of SARS-CoV-2 3CL^pro^ as well as machine intelligence[18–23]. Highly potent SARS-CoV-2 3CL^pro^ inhibitors with diverse chemical structures need further exploration.

Traditional Chinese medicine (TCM) herbs and formulae have long been used in treating viral diseases. Some of them have been clinically tested to treat COVID-19[24]. Scutellariae radix (Huangqin in Chinese), the root of *Scutellaria baicalensis* Georgi, has been widely used in TCM for heat clearing, fire purging, detoxification and hemostasis. Huangqin is officially recorded in Chinese Pharmacopoeia (2015 Edition)[25] and European Pharmacopoeia (10^th^ Edition)[26]. Its anti-tumor, antiviral, anti-microbial and anti-inflammatory activities have been reported[27]. Remarkably, the extracts of *S. baicalensis* have exhibited broad spectrum anti-viral activities, including ZIKA[28], H1N1[29], HIV[30] and DENV[31]. In addition, a multicenter, retrospective analysis demonstrated that *S. baicaleinsis* exhibits more potent antiviral effects and higher clinical efficacy than ribavirin for the treatment of hand, foot and mouth disease[32]. Several *S. baicalensis* derived mixtures or pure compounds have been approved as antiviral drugs, such as Baicalein capsule (to treat hepatitis) and Huangqin tablet (to treat upper respiratory infection) in China. Most of the *S. baicaleinsis* ingredients are flavonoids[33]. Flavonoids from other plants were also reported to mildly inhibit SARS and MERS-CoV 3CL^pro^ [34, 35]. Here we studied the anti-SARS-CoV-2 activity of *S. baicalensis* and its ingredients. We found that the ethanol extract of *S. baicalensis* inhibits SARS-CoV-2 3CL^pro^ activity and the most active ingredient baicalein exhibits an IC50 of 0.39 μM. In addition, the ethanol extract of *S. baicalensis* effectively inhibits the replication of SARS-CoV-2 in cell assay. We also identified four baicalein analogue compounds from other herbs that inhibit SARS-CoV-2 3CL^pro^ activity at microM concentration.

## Results and Discussion

### The ethanol extract of *S. baicalensis* strongly inhibits SARS-CoV-2 3CL^pro^

We prepared the 70% ethanol extract of *S. baicalensis* and tested its inhibitory activity against SARS-CoV-2 3CL^pro^. We expressed SARS-CoV-2 3CL^pro^ and performed activity assay using a peptide substrate (Thr-Ser-Ala-Val-Leu-Gln-pNA) according to the published procedure of SARS-CoV 3CL^pro^ assay[11, 36]. The inhibitory ratio of *S. baicalensis* extract at different concentrations on SARS-CoV-2 3CL^pro^ activity were shown in Figure 1A. The crude extract exhibits significant inhibitory effect with an IC_50_ of 8.5 μg/ml, suggesting that *S. baicalensis* contains candidate inhibitory ingredients against SARS-CoV-2 3CL^pro^.

**Figure 1.**
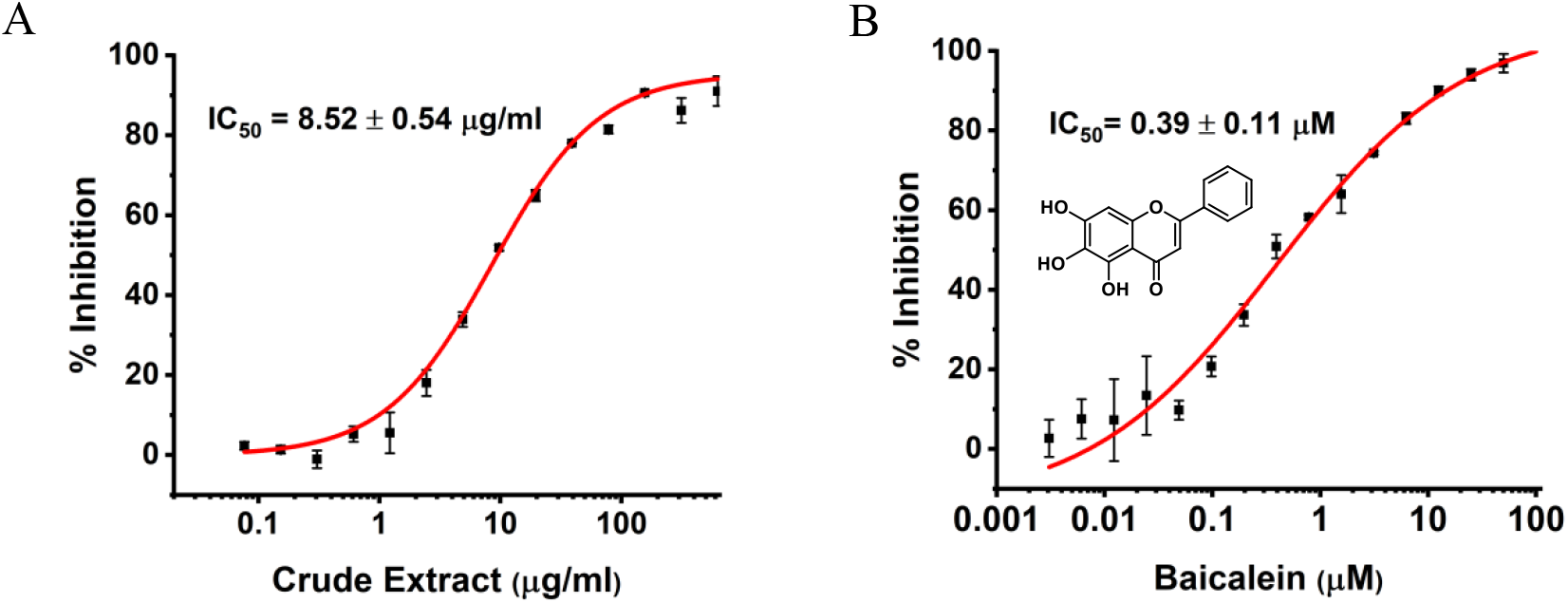
The *in vitro* anti-SARS-CoV-2 3CL^pro^ activity of *S. baicalensis* ethanol extract (A) and baicalein (B).

### Baicalein is the major active ingredient in *S. baicalensis* that inhibits SARS-CoV-2 3CL^pro^

We tested the inhibitory activity of four major ingredients from *S. baicalensis*: baicalein, baicalin, wogonin and wogonoside *in vitro*. Baicalein showed the most potent anti-SARS-CoV-2 3CL^pro^ activity with an IC_50_ of 0.39 μM (Figure 1B and Table 1). Baicalin inhibited SARS-CoV-2 3CL^pro^ activity for about 41% at 50 μM, while wogonin and wogonoside were not active at this concentration.

**Table 1.**
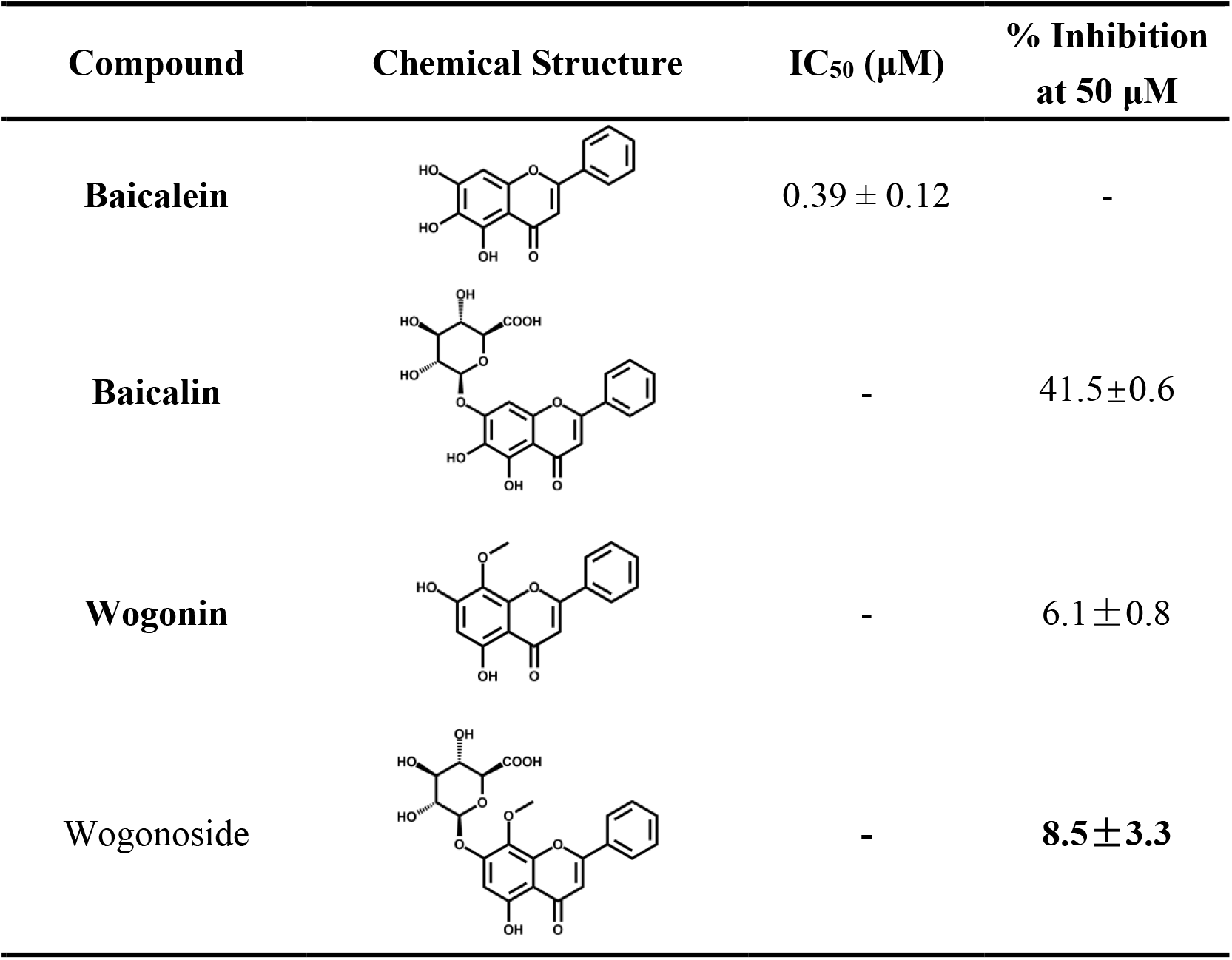
The SARS-CoV-2 3CL^pro^ inhibition activity of four major flavones derived from *S. baicalensis*.

| Compound | Chemical Structure | IC<sub>50</sub> (μM) | % Inhibition at 50 μM |
| --- | --- | --- | --- |
| Baicalein |  | 0.39 ± 0.12 | - |
| Baicalin |  | - | 41.5±0.6 |
| Wogonin |  | - | 6.1±0.8 |
| Wogonoside |  | - | 8.5±3.3 |

**Table 2.**
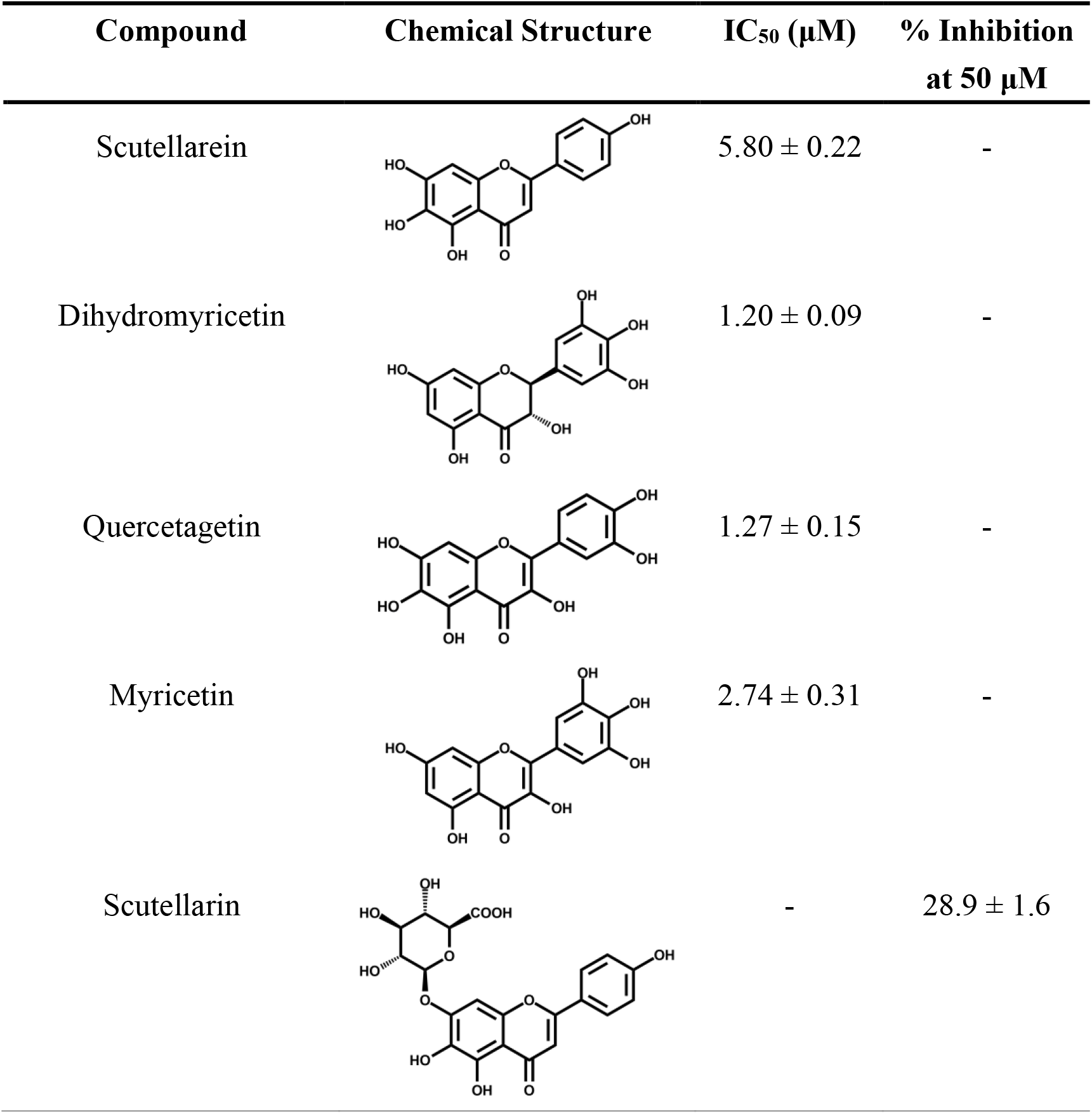
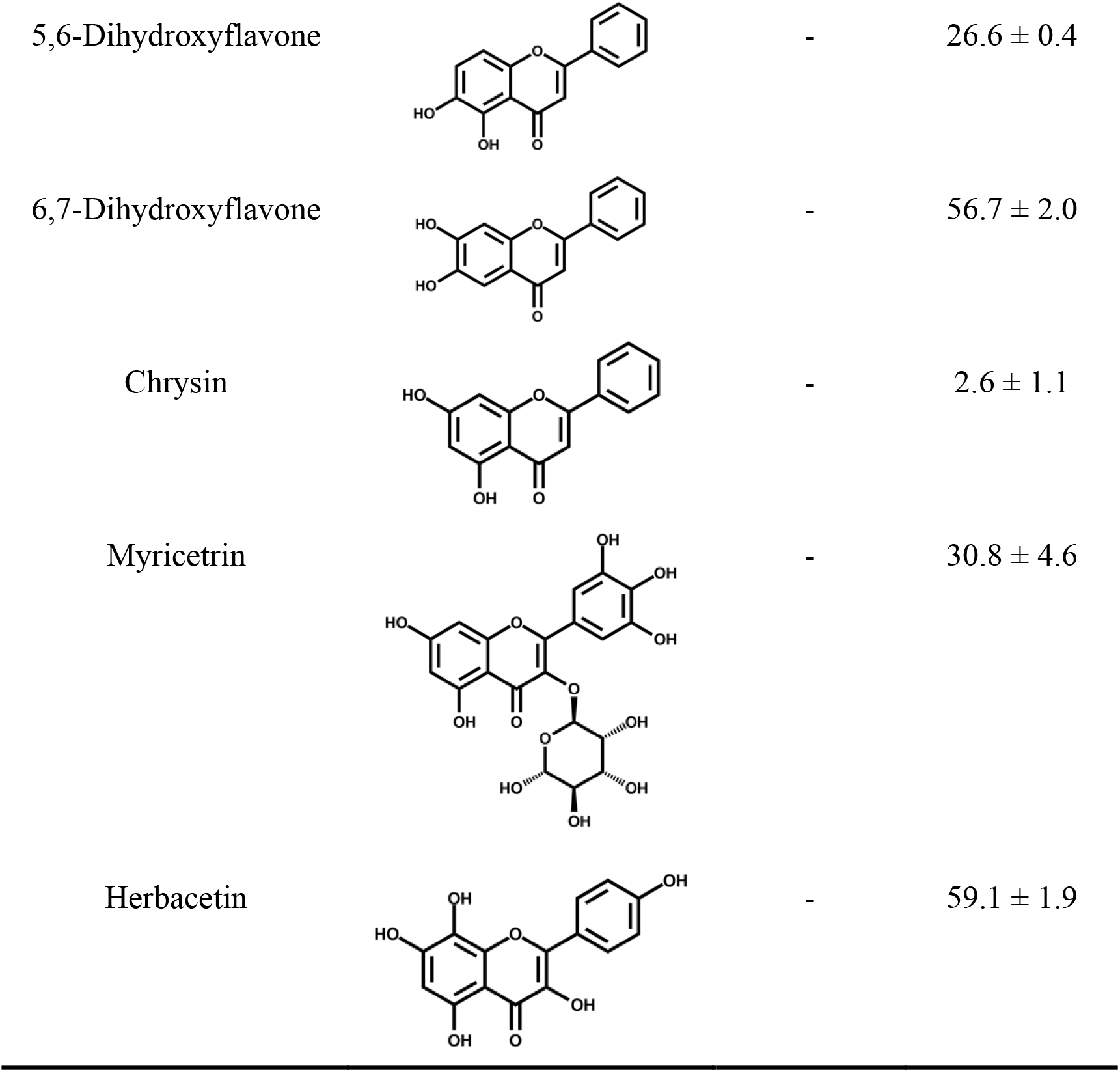
The anti-SARS-CoV-2 3CL^pro^ activity of baicalein analogue flavonoids.

| Compound | Chemical Structure | IC<sub>50</sub> (μM) | % Inhibition at 50 μM |
| --- | --- | --- | --- |
| Scutellarein |  | 5.80 ± 0.22 | - |
| Dihydromyricetin |  | 1.20 ± 0.09 | - |
| Quercetagetin |  | 1.27 ± 0.15 | - |
| Myricetin |  | 2.74 ± 0.31 | - |
| Scutellarin |  | - | 28.9 ± 1.6 |
| 5,6-Dihydroxyflavone | | - | $26.6 \pm 0.4$ |
| 6,7-Dihydroxyflavone | | - | $56.7 \pm 2.0$ |
| Chrysin | | - | $2.6 \pm 1.1$ |
| Myricetrin | | - | $30.8 \pm 4.6$ |
| Herbacetin | | - | $59.1 \pm 1.9$ |

We performed molecular docking to understand the inhibitory activity of *S. baicalensis* ingredients. In the docking model, baicalein binds well in the substrate binding site of SARS-CoV-2 3CL^pro^ with its 6-OH and 7-OH forming hydrogen bond interactions with the carbonyl group of L141 and the backbone amide group of G143, respectively (Figure 2A). In addition, the carbonyl group of baicalein is hydrogen bonded with the backbone amide group of E166. The catalytic residues H41 and C145 are well covered by baicalein, accounting for its inhibitory effect. As the 7-OH in baicalin is in close contact with the protein, there may not be enough space for glycosyl modification, explaining the low activity of baicalin. As for wogonin, the absence of 6-OH together with its additional 8-methoxyl group alters the binding orientation and weakens the binding strength (Figure 2B). Hydrogen bond is observed between its 5-OH and the backbone carbonyl group of L141, while the interaction with E166 by its 8-methoxy group is weaker than that formed by the carbonyl group in baicalein.

**Figure 2.**
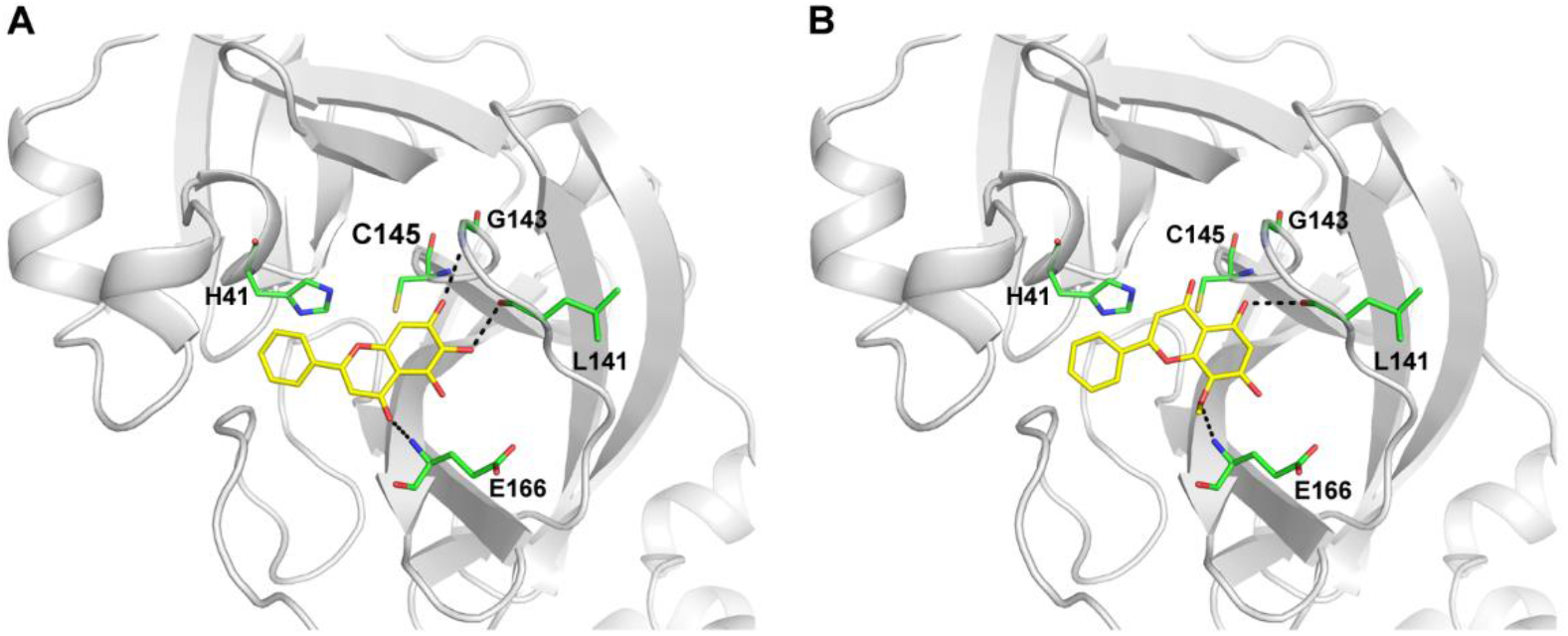
The interactions between SARS-CoV-2 3CL^pro^ and *S. Baicalensis* ingredients baicalein (A) and wogonin (B) in the docking models. The overall structure and key residues of SARS-CoV-2 3CL^pro^ are shown as grey cartoon and green sticks, respectively. *S. Baicalensis* ingredients are displayed as yellow sticks.

### *S. baicalensis* extract and baicalein inhibit the replication of SARS-CoV-2 in Vero cells

We tested the antiviral activity of *S. baicalensis* ethanol extract and baicalein against SARS-CoV-2 using RT-qPCR. Vero cells were pre-treated with the extract or baicalein for 1h, followed by virus infection for 2h. Virus input was then washed out and the cells were treated with medium containing the extract or baicalein. Viral RNA was extracted from the supernatant of the infected cells and quantified by RT-PCR. The *S. baicalensis* ethanol extract significantly reduced the growth of the virus with an EC_50_ of 0.74 μg/ml with low cytotoxicity (SI > 675.68, Figure 3A). Baicalein inhibits the replication of SARS-CoV2 with an EC_50_ of about 17.6 μM and SI > 2.8 (Figure 3B). The high activity of *S. baicalensis* crude extract in the antiviral assay implies it may also interact with other viral or host targets in addition to SARS-CoV-2 3CL^pro^ inhibition, which can be further explored in the future.

**Figure 3.**
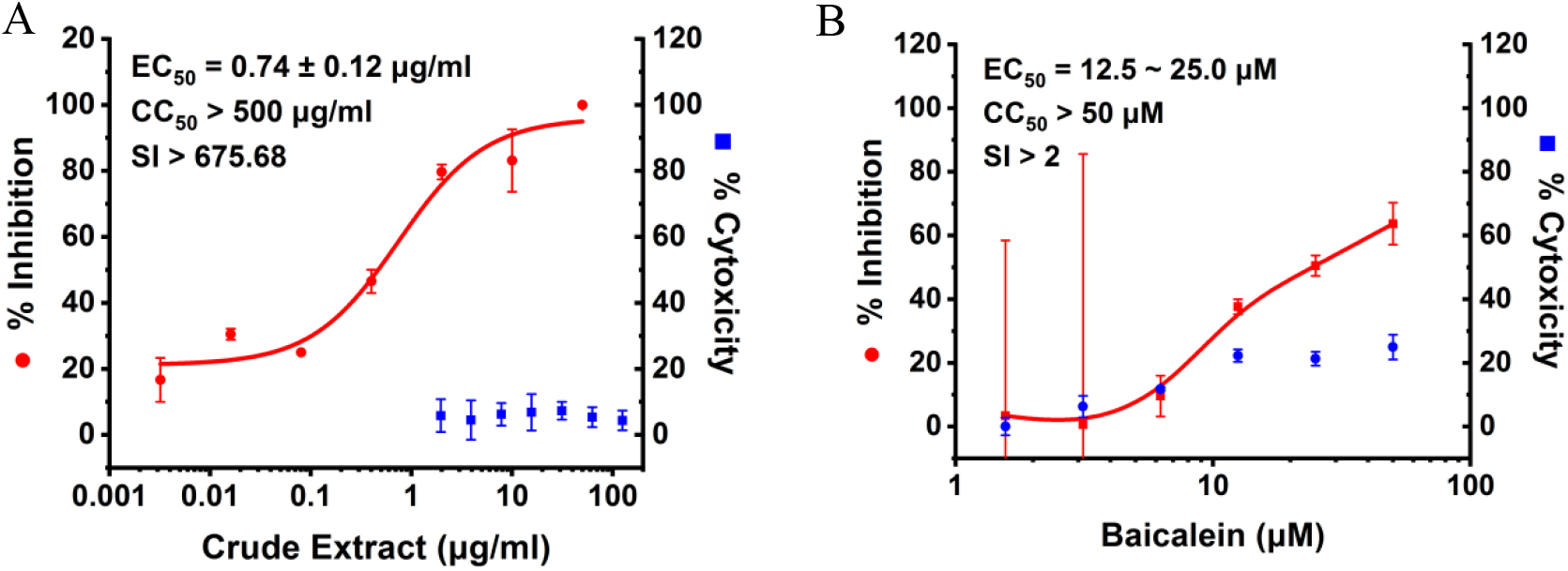
The antiviral activity of *S. baicalensis* extract (A) and baicalein (B) against SARS-CoV-2 in Vero cells.

**Figure 3.**
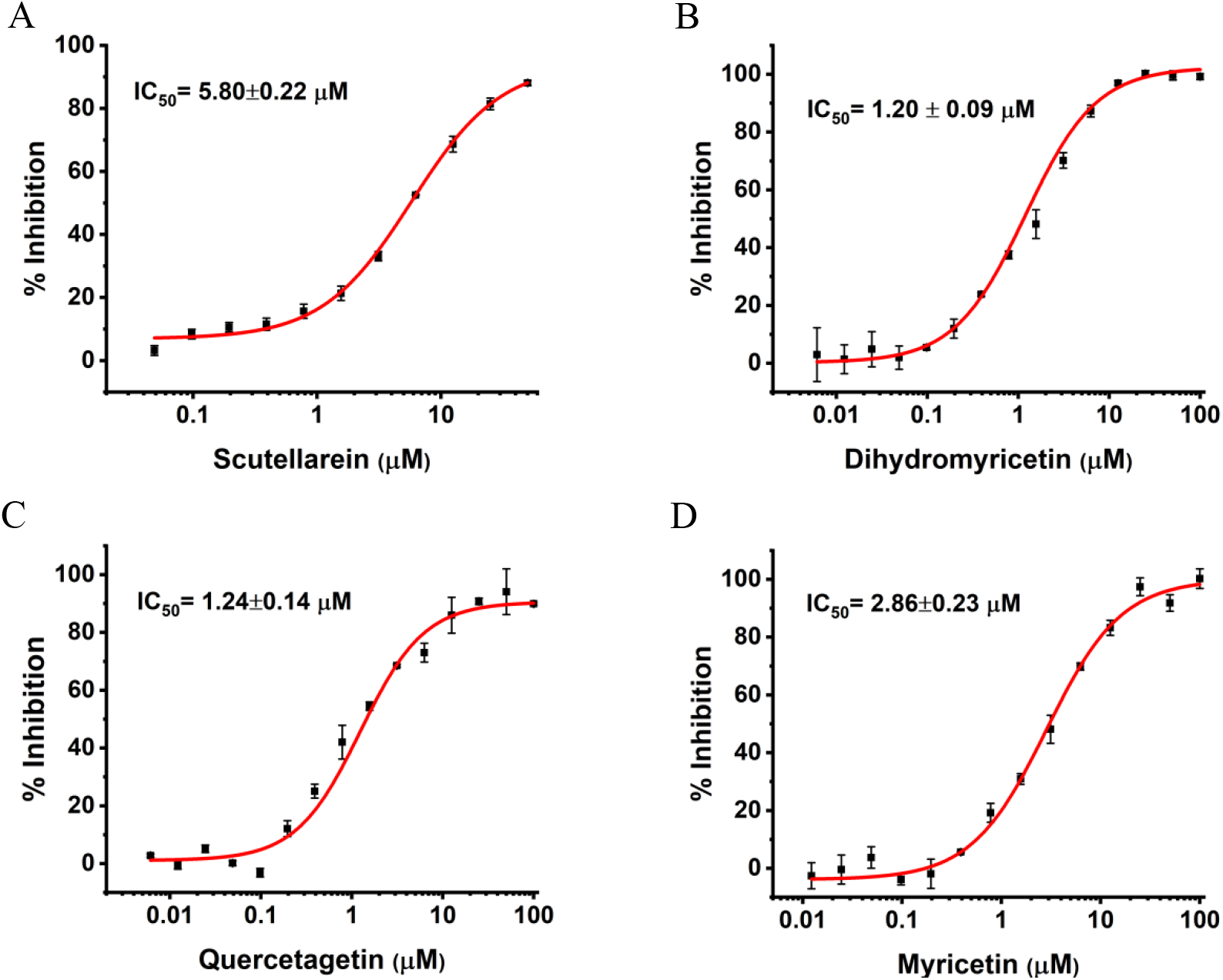
The SARS-CoV-2 3CL^pro^ inhibition activity of (A) scutellarein, (B) dihydromyricetin, (C) quercetagetin and (D) myricetin.

### Searching for baicalein analogues that inhibit SARS-CoV-2 3CL^pro^

We searched for baicalein analogues from available flavonoid suppliers and selected 8 flavonoids and 2 glycosides for experimental testing. Four flavonoid compounds were found to be potent SARS-CoV-2 3CL^pro^ inhibitors. Among them, scutellarein is mainly distributed in genus *Scutellaria* and Erigerontis herba (Dengzhanxixin or Dengzhanhua in Chinese) in its glucuronide form, scutellarin. Scutellarin has long been used in cardiovascular disease treatment for its ability to improve cerebral blood supply[37]. Scutellarein inhibits SARS-CoV-2 3CL^pro^ with an IC_50_ value of 5.8 μM, while scutellarin showed mild inhibitory activity at 50 μM concentration. The other three flavonoid compounds, dihydromyricetin, quercetagetin and myricetin derived from *Ampelopsis japonica* (Bailian in Chinese), *Eriocaulon buergerianum* (Gujingcao in Chinese) and *Polygoni avicularis* (Bianxu in Chinese) respectively, inhibit SARS-CoV-2 3CL^pro^ with IC_50_ values of 1.20, 1.24 and 2.86 μM. Interestingly, scutellarein and myricetin were reported to inhibit the SARS-CoV indicating their potential as multi-target anti-SARS-CoV-2 agents[38].

For all the active flavonoid compounds that we found, the introduction of glycosyl group, as in the case of baicalein and baicalin, decreased the inhibition activity, probably due to the steric hindrance of the glycosyl group, which is also true for scutellarein/scutellarin, and myricetin/myricetrin. As glycosides and their corresponding aglycones are often interchangeable *in vivo*, for instance, baicalin was reported to be metabolized to baicalein in intestine[39], while baicalein can be transformed to baicalin by hepatic metabolism[40], we expect that both the flavonoid form of the active compounds and their glycoside form will function *in vivo*. We suggest that these compounds can be further optimized or used to search for other TCM herbs containing these compounds or substructures for the treatment of COVID-19.

## Material and methods

*S. Baicalensis* were purchased from Tong Ren Tang Technologies Co. Ltd. Baicalein and compounds not listed below were from J&K Scientific. 5,6-dihydroxyflavone was purchased from Alfa Aesar. 6,7-dihydroxyflavone was synthesized by Shanghai Yuanye Biotechnology Co., Ltd. Myricetin, quercetagetin and herbacetin were purchased from MCE. Dihydromyricetin and myricetrin were purchased from Targetmol.

### Construction of plasmid SARS-CoV-2 pET 3CL-21x, protein expression and purification

The DNA of SARS-CoV-2 3CL^pro^ (referred to GenBank, accession number MN908947) was synthesized (Hienzyme Biotech) and amplified by PCR using primers n3CLP-Nhe (5’-CATGGCTAGCGGTTTTAGAAAAATGGCATTCCC-3’) and n3CLP-Xho (5’-CACTCTCGAGTTGGAAAGTAACACCTGAGC-3’). The PCR product was digested with *Nhe* I/*Xho* I and cloned into the pET 21a DNA as reported previously [41]. The resulting SARS-CoV-2 pET 3CL-21x plasmid encodes a 35 064 Da SARS-CoV-2 3CL^pro^ with a C-terminal 6xHis-tag. The SARS-CoV-2 pET 3CL-21x plasmid was further transformed to *E. coli* BL21<DE3> for protein expression as reported [41]. The recombinant protein was purified through a nickel-nitrilotriacetic acid column (GE Healthcare) and subsequently loaded on a gel filtration column Sephacryl S-200 HR (GE Healthcare) for further purification as previously described [42].

### Enzyme inhibition assay

A colorimetric substrate Thr-Ser-Ala-Val-Leu-Gln-pNA (GL Biochemistry Ltd) and assay buffer (40 mM PBS, 100 mM NaCl, 1 mM EDTA, 0.1% Triton 100, pH 7.3) was used for the inhibition assay. Stock solutions of the inhibitor were prepared with 100% DMSO. The 100 μl reaction systems in assay buffer contain 0.5 μM protease and 5% DMSO or inhibitor to the final concentration. Firstly, dilute SARS-CoV-2 3CL^pro^ with assay buffer to the desired concentration. 5 μl DMSO or inhibitor at various concentrations was pre-incubated with 85 μl dilute ed SARS-CoV-2 3CL^pro^ for 30 min at room temperature. And then add 10 μl 2 mM substrate Thr-Ser-Ala-Val-Leu-Gln-pNA (dissolved in water) into above system to final concentration of 200 μM to initiate the reaction. Increase in absorbance at 390 nm was recorded for 20 min at interval of 30 s with a kinetics mode program using a plate reader (Synergy, Biotek). The percent of inhibition was calculated by V_i_/V_0_, where V_0_ and V_i_ represent the mean reaction rate of the enzyme incubated with DMSO or compounds. IC_50_ was fitted with Hill1 function.

### Molecular docking

The structure of SARS-CoV-2 3CL^pro^ (PDB ID 6LU7)[14] and *S. baicalensis* components were prepared using Protein Preparation Wizard and LigPrep module, respectively. Then, the binding site was defined as a 20*20*20 Å^3^ cubic box centered to the centroid of C145. After that, molecular docking was performed using Glide. Extra precision (XP) and flexible ligand sampling were adopted. Post-docking minimization was performed to further refine the docking results. All the above mentioned modules were implemented in Schrödniger version 2015-4 (Schröidnger software suite, L. L. C. New York, NY (2015).)

### Cell culture and virus

Vero cell line (ATCC, CCL-81) was cultured at 37 °C in Dulbecco’s modified Eagle’s medium (DMEM, Gibco, Grand Island, USA) supplemented with 10% fetal bovine serum (FBS, Gibco) in the atmosphere with 5% CO_2_. Cells were digested with 0.25% trypsin and uniformly seeded in 96-well plates with a density of 2×10^4^ cells/well prior infection or drug feeding. The virus (C-Tan-nCoV Wuhan strain 01) used is a SARS-COV-2 clinically isolated virus strain. These viruses were propagated in Vero cells.

### Antiviral activity Assay

The cytotoxicity of *S. baicalensis* extract and baicalein on Vero cells were determined by CCK8 assays (DOJINDO, Japan). We then evaluated the antiviral efficiency of *S. Baicalensis* extract and baicalein against SARS-COV-2 (C-Tan-nCoV Wuhan strain 01) virus *in vitro*. Cells were seeded into 96-well plates at a density of 2×10^4^ cells/well and then grown for 24 hours. Cells were pre-treated with indicated concentrations of *S. Baicalensis* extract or baicalein for 1 h, and the virus (MOI of 0.01, 200 PFU/well) was subsequently added to allow infection for 2 h at 37℃.Virus input was washed with DMEM and then the cells were treated with medium contained drugs at various concentrations for 48h. The supernatant was collected and the RNA was extracted and analyzed by relative quantification using RT-PCR as in the previous study[3,43].

### RNA extraction and RT-qPCR

Viral RNA was extracted from 100 μL supernatant of infected cells using the automated nucleic acid extraction system (TIANLONG, China), following the manufacturer’s recommendations. SARS-COV-2 virus detection was performed using the One Step PrimeScript RT-PCR kit (TaKaRa, Japan) on the LightCycler 480 Real-Time PCR system (Roche, Rotkreuz, Switzerland). ORF 1ab was amplified from cDNA and cloned into MS2-nCoV-ORF1ab and used as the plasmid standard after its identity was confirmed by sequencing. A standard curve was generated by determination of copy numbers from serially dilutions (10^3^−10^9^ copies) of plasmid. The following primers used for quantitative PCR were 1ab-F: 5ʹ-AGAAGATTGGTTAGATGATGATAGT-3ʹ; 1ab-R: 5ʹ-TTCCATCTCTAATTGAGGTTGAACC-3ʹ; and probe 5ʹ-FAM-TCCTCACTGCCGTCTTGTTG ACCA-BHQ1-3ʹ. The individual EC_50_ values were calculated by the Origin 2018 software.

## Acknowledgements

This work was supported in part by the Ministry of Science and Technology of China (2016YFA0502303, 2016YFD0500301), the National Natural Science Foundation of China (21633001) and Peking University Special Fund for COVID-19.

## Notes

### Competing Interest Statement

The authors have declared no competing interest.

